# The VCBS superfamily, a diverse group of β-propellers that includes tachylectin and integrins

**DOI:** 10.1101/2020.10.09.333302

**Authors:** Joana Pereira, Andrei N. Lupas

**Affiliations:** Department of Protein Evolution, Max-Planck-Institute for Developmental Biology, Tübingen 72076, Germany

## Abstract

β-Propellers are found in great variety across all kingdoms of life. They assume many cellular roles, primarily as scaffolds for macromolecular interactions and catalysis. Despite their diversity, most β-propeller families clearly originated by amplification from the same ancient peptide - the β-propeller blade. In cluster analyses, β-propellers of the WD40 superfamily always formed the largest group, to which some important families, such as the α-integrin, Asp-box, and glycoside hydrolase β-propellers connected weakly. Motivated by the dramatic growth of sequence databases we revisited these connections, with a special focus on VCBS-like β-propellers, which have not been analysed for their evolutionary relationships so far. We found that they form a supercluster with integrin-like β-propellers and tachylectins, clearly delimited from the superclusters formed by WD40 and Asp-Box β-propeller. Connections between the three superclusters are made mainly through PQQ-like β-propeller. Our results present a new, greatly expanded view of the β-propeller classification landscape.

## Introduction

Proteins with a β-propeller domain are found in all kingdoms of life (Fig. S1c). They are involved in diverse biological processes, from adhesion to transcription regulation (Fülöp and Jones 1999; Pons et al. 2003; Guruprasad and Dhamayanthi 2004; Chen et al. 2011). In them, the β-propeller acts mostly as a recognition site for different biomolecules, but may also carry catalytic activity. These repetitive domains (Andrade et al. 2001; Söding and Lupas 2003) adopt a toroid fold, where between 4 and 12 (Fig. S1d) copies of a widespread supersecondary structure, the 4-stranded β-meander, are arranged radially around a central channel (Fig. S1a,b). These repeats, whose strands are labelled A to D (Figs. 2b, 3 and S1b), are called ‘blades’ and the toroids they form correspondingly ‘propellers’. Blades carry specific sequence motifs which allow the classification of cognate β-propellers into a hierarchy of families and superfamilies (Fülöp and Jones 1999; Pons et al. 2003; Guruprasad and Dhamayanthi 2004; Chaudhuri et al. 2008; Chen et al. 2011).

Despite their wide sequence diversity (Fig. S1e,f), most β-propeller families are related to each other and emerged by independent amplification from a set of homologous ancestral blades, in a process that is still visibly ongoing (Chaudhuri et al. 2008; Kopec and Lupas 2013; Dunin-Horkawicz et al. 2014; Alva et al. 2015; Afanasieva et al. 2019). Classification studies suggested that most β-propeller families form a supercluster centred on WD40 β-propellers, a large superfamily characterised by a Trp-Asp motif at the end of strand C (in position 40) (Fig. 2b). Proteins assigned to this supercluster in previous studies included the β-subunits of G-proteins, the low density lipoprotein (LDL) receptors, protein kinase PknD and the tachyletin-2 family, which comprises eukaryotic lectins involved in the innate immunity of cnidarians and crustaceans (Neer et al. 1994; Beisel et al. 1999; Chaudhuri et al. 2008; Hayes et al. 2010; Kopec and Lupas 2013). Some peripheral groups connected weakly to this supercluster (Chaudhuri et al. 2008; Kopec and Lupas 2013), such as the β-propeller domain of α-integrins, characterised by a Ca^2+^-binding DxDxDG motif in the loop connecting strands A and B (loop AB) and an FG-GAP/Cage motif, which is contiguous in space but not sequence, covering the N-terminal end of strand A and the C-terminal end of strand B (Rigden and Galperin 2004; Chouhan et al. 2011). This connection was proposed to be weakly mediated by Asp-Box β-propellers, most of whose members are characterized by a SxDxGxTW motif in the loop connecting strands C and D (loop CD) (Quistgaard and Thirup 2009).

Missing from these studies were β-propellers of the *Vibrio, Colwellia, Bradyrhizobium,* and *Shewanella* (VCBS) family (Pfam: PF1351), a poorly described group that has hitherto not been analysed systematically for its evolutionary relationships. VCBS encompasses the β-propellers in aldos-2-ulose dehydratases (AUDH) (Claesson et al. 2012), ABC toxin component B (TcB) (Meusch et al. 2014), fungal PVL lectins (Cioci et al. 2006), and a variety of hypothetical archaeal toxins (Makarova et al. 2019). As PVLs carry a conserved Ca^2+^-binding DxDxDG motif in loop AB, their similarity to integrin-like β-propellers has been conjectured (Cioci et al. 2006), but their mode of carbohydrate recognition appears to be more similar to that of tachylectin-2 (Beisel et al. 1999; Cioci et al. 2006). In order to obtain further insight into this group and locate it within the β-propeller landscape, we performed a survey of VCBS-like β-propellers and their relationship to integrin-like, Asp-Box, tachylectin and WD40 β-propellers.

## Results and Discussion

PSI-BLAST searches with 13 β-propellers of known structure, chosen to represent the families described above (Table S1), yielded a total of 5996 sequences from bacteria, archaea and eukaryotes (see Methods). When clustered by pairwise similarity (Fig. 1), these sequences form three superclusters organized around cores of WD40, Asp-Box and VCBS-like β-propellers, respectively. The WD40 and Asp-Box superclusters were expected, based on previous analyses (Chaudhuri et al. 2008; Kopec and Lupas 2013), but we were struck by the clear grouping of the other β-propeller families into a third supercluster, centered on VCBS and clearly delimited from the other two.

**Figure 1.**
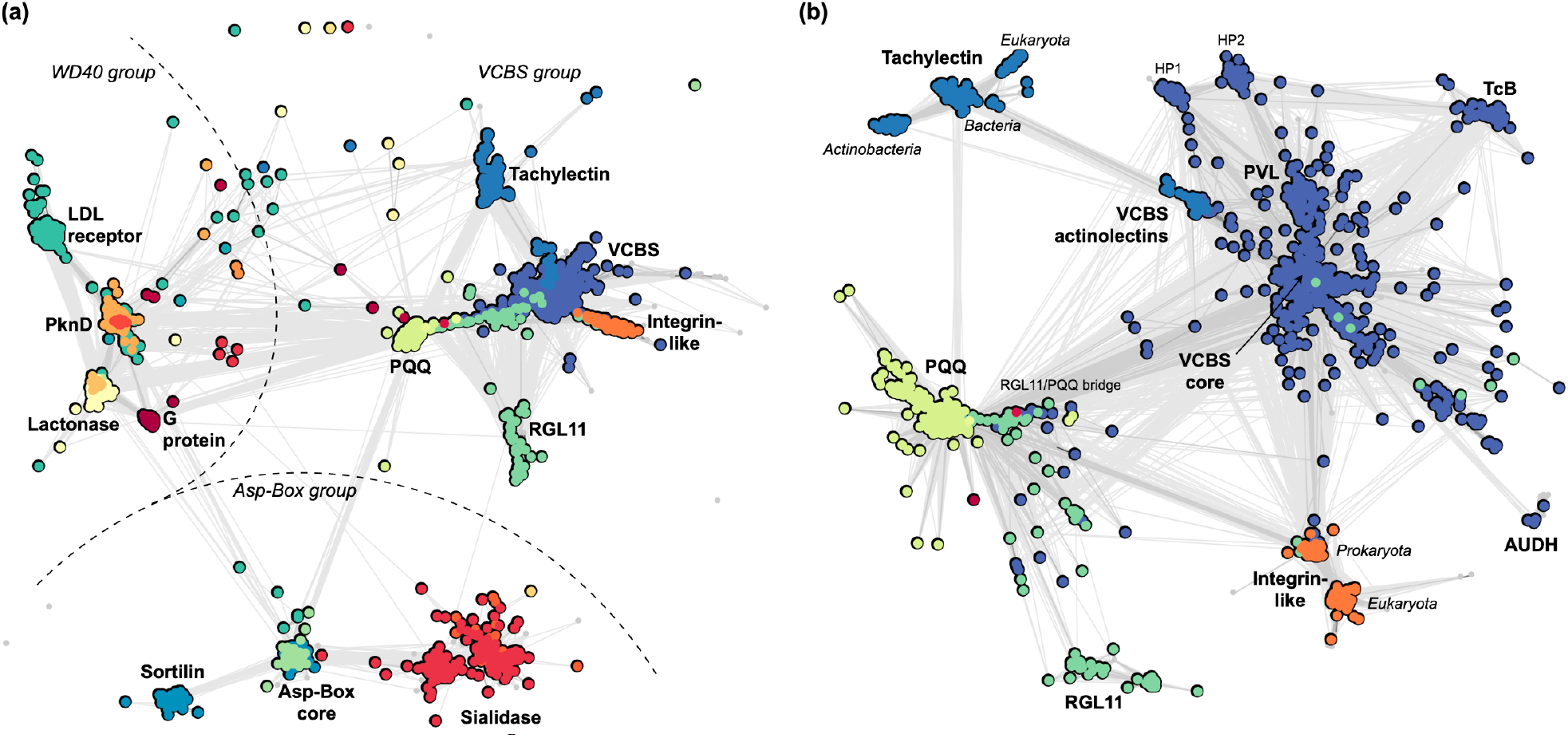
Classification landscape of representative β-propeller families, coloured based on the family name (f-name) of the best match in HMM searches against ECOD. (a) Cluster map of all 5996 sequences collected. Clustering was carried out with CLANS in 2D until equilibrium at a BLASTp p-value of 10^−5^, with connections represent similarities at this p-value (the darker, the more similar). Different regions of the map are annotated with the name of the sequences within the corresponding cluster or, when a cluster encompasses multiple families, by the β-propeller family as in ECOD and Pfam. (b) Cluster map of the 2662 sequences in the VCBS supercluster. Clustering was carried out as in (a) but a BLASTp p-value of 10^−18^, in order to expand it and uncover its internal structure. Connections are shown at a BLASTp p-value of 10^−10^. Multiple colours within the same cluster correspond to sequences that match multiple close β-propeller families. HP stands for ‘hypothetical propeller’.

The core of the VCBS supercluster comprises prokaryotic β-propellers from diverse hypothetical protein families (Fig. S2), which carry a signal sequence and may contain several β-propeller domains, accompanied by domains associated with biomolecular interactions (mostly immunoglobulin-like domains, but also armadillo repeats and jelly-roll-like lectins, Fig. S2). The VCBS core group is connected to a large periphery of VCBS-like families, including PVL, TcB and AUDH, as well as to diverse hypothetical β-propellers, which have hitherto remained unstudied (Figs. 1b and S2). β-Propeller families in this periphery are found in a variety of hypothetical proteins, whose domain composition suggests an involvement in biomolecular interactions and catalysis (Fig. S2a). The most peripheral families that still connect directly to the VCBS core are the integrin-like β-propellers and the bacterial RGL11 family (rhamnogalacturonan lyase YesX, ECOD: 001396995). Two other important β-propeller families complete the VCBS supercluster, comprising tachylectins and PQQ β-propellers, respectively. These connect to each other, and also to the VCBS core via RGL11, in the case of PQQ, and a β-propeller family we have named VCBS actinolectins, in the case of tachylectins.

We chose the name “VCBS actinolectins” given their exclusive occurrence in actinobacteria and evolutionary connection to tachylectins (Figs.1b and S2), but no member of this family has as yet been characterized. These β-propellers are found in proteins that carry a signal sequence and either consist of the single β-propeller domain or of the β-propeller preceded by a TIM barrel (Figs. S2a). Their connection to the tachylectin cluster is mediated by a core of bacterial tachylectin-like sequences, which are found in secreted proteins often containing additional domains involved in catalysis. Two groups radiate from this core, the eukaryotic tachylectins-2 and a second family of actinobacterial β-propellers, both of which are comprised of secreted proteins consisting of the β-propeller domain alone. The identification of these multiple tachylectin-like families was a striking result as tachylectin β-propellers have been considered for long time as near-orphans and have so far only been reported in eukaryotes (Beisel et al. 1999; Hayes et al. 2010; Smock et al. 2016).

HMM comparisons highlight the sequence motifs behind the connections described here (Fig. 2). The most prominent motif is the aspartate-rich DxDxDG sequence of loop AB (Figs. 2b and 3) (Rigden and Galperin 2004; Cioci et al. 2006; Chouhan et al. 2011). While in PVL and α-integrin, this loop binds Ca^2+^ (Fig. 3b), in other members it may recognise also other metal cations (Rigden and Galperin 2004; Chouhan et al. 2011; Claesson et al. 2012; Meusch et al. 2014). Also conspicuous are two non-contiguous, highly conserved residues of loop CD, G and W (Fig. 2b). Their functional role is uncertain, but in integrin-like β-propellers the G coordinates a water molecule involved in Ca^2+^ binding (Chouhan et al. 2011), and in tachylectin-2 the W anchors a short α-helix involved in forming the sugar-binding pocket (Fig. 3). A fourth prominent motif is a GW in loop DA’ (the loop that connects strand D from one blade to strand A of the next) (Figs. 2b and 3a,c), which in tachylectin-2 and PVL is involved in forming the sugar-binding pocket (Fig. S3) (Kawabata and Tsuda 2002; Cioci et al. 2006).

**Figure 2.**
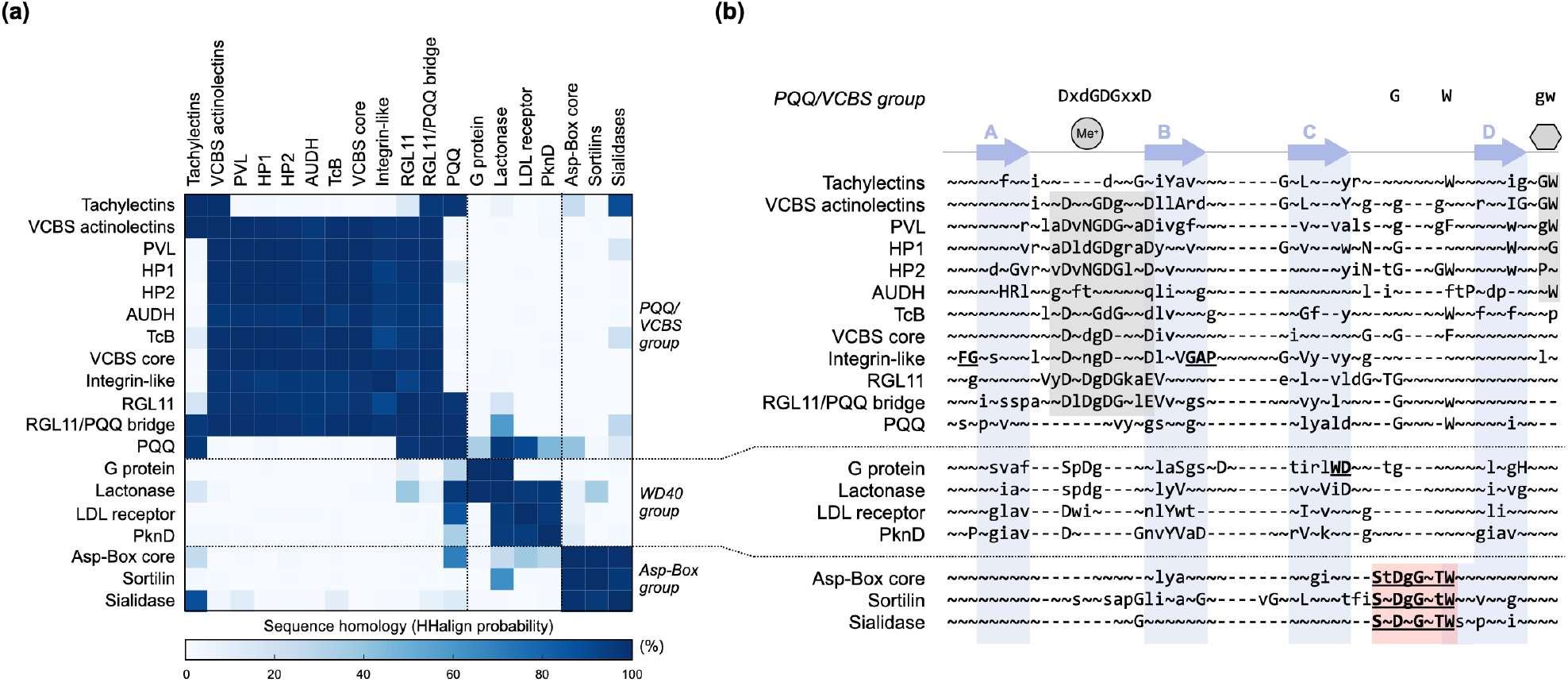
HMM comparison of β-propeller groups. (a) Sequence homology matrix of β-propeller groups selected from the cluster maps, as measured by the probability of the alignment of full-length HMM profiles with HHalign. (b) Multiple alignment of the HMM consensus sequences, focused on single-bladed regions. Sequence motifs common to the VCBS supercluster are highlighted in grey and summarised on top. Their function in members of known structure is depicted: a grey circle with Me^+^ represents ‘metal binding’ and a grey hexagon ‘sugar binding’. The Asp-Box motif is highlighted in light red. Arrows depict the four strands of blade and are named accordingly. This annotation was carried out based on the known structures of families shown, but represent only a consensus as, due to structural deviations or especial structural features, the specific start and end of these strands may be shifted.

**Figure 3.**
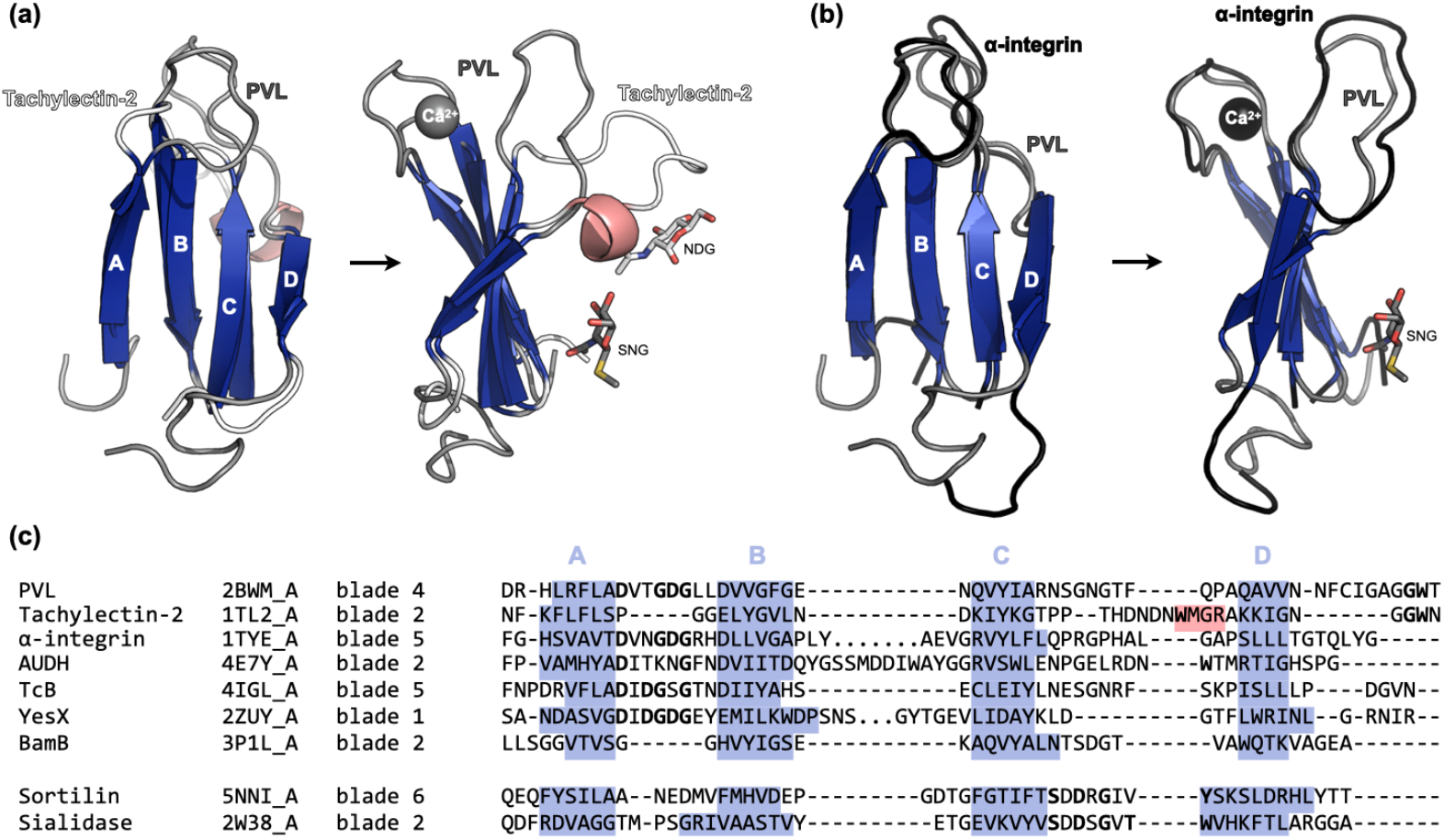
Structure-based alignment of representative blades of the VCBS-PQQ and Asp-Box superclusters. (a-b) Structural superposition of the 4^th^ blade of fungal PVL lectin (pdbID: 2BWM_A) to (a) the 2^nd^ blade of the tachylectin-2 β-propeller (pdbID: 1TL2_A) and (b) the 5^th^ blade in the α-integrin β-propeller (pdbID: 1TYE_A). Ligands are highlighted, coloured according to the parental protein. SNG: methyl 2-acetamido-2-deoxy-1-seleno-beta-D-glucopyranoside; NDG: 2-acetamido-2-deoxy-alpha-D-glucopyranose. (c) Structure-based sequence alignment of the 4^th^ blade of fungal PVL lectin to individual representative blades. The pdbID as well as the corresponding blade indices are shown. Residues in stranded regions are highlighted in blue and those in helical regions in light red.

While widely represented in the families of the VCBS supercluster, none of these motifs is universal. Thus, for example, the aspartate-rich motif of loop AB is not found in tachylectin-like and PQQ β-propellers. These are connected to other families in the supercluster by the sequence of loop CD and, in the case of tachylectin-like β-propellers, by the GW motif of loop DA’.

## Conclusions

Our results confirm the relationship conjectured between fungal PVL lectins, tachylectin-2 and integrin-like β-propellers (Cioci et al. 2006). We find that all three of these eukaryotic protein families are satellites of larger prokaryotic clusters, from which they are presumably descended. Jointly with these, they are part of a supercluster of β-propeller families, centred on the large group of prokaryotic VCBS β-propellers. This supercluster had not been recognised in previous studies (Chaudhuri et al. 2008; Kopec and Lupas 2013) because most relevant proteins could not be included, primarily due to the lack of relevant sequences of known structure.

We believe two factors were essential in our ability to resolve the evolutionary connections between the main β-propeller groups. The first is the presence of members of the VCBS superfamily, which revealed their intermediate position between integrin-like and PQQ β-propellers, providing a context for the weak links previously observed between integrin-like and Asp-Box β-propellers. The second was the collection of a substantial number of tachylectin-like sequences. Given the structural approach of previous studies (Chaudhuri et al. 2008; Kopec and Lupas 2013), these encompassed only the one tachylectin-like sequence found in PDB, which clustered in the WD40 supercluster. In our study, more than 140 tachylectin-like sequences were collected, including sequence intermediates essential for the establishment of evolutionary links. Many of these sequences are of bacterial origin and resulted from metagenomic studies, highlighting the importance of such efforts for the better understanding of protein evolution paths and the structure of the β-propeller sequence space.

## Materials and Methods

Thirteen β-propeller representatives of known structure (Table S1) were used as queries for sequence searches with PSI-BLAST (Altschul 1997). Searches were carried out with the *nr* database filtered to a maximum sequence identity of 30% (*nr30*, as of May 2020) (Zimmermann et al. 2018). Tachylectins were searched on the *nr* database filtered to a maximum sequence identity of 50%. Matches covering more than 80% of the corresponding query were collected after 2 rounds and filtered to a maximum sequence identity of 50% with CD-HIT (Li and Godzik 2006). The final sequences were assigned an ECOD family with HHsearches against a database of HMM profiles for the ECOD database filtered to 70% maximum sequence identity (HHpred ECOD70 database as of March 2020) (Zimmermann et al. 2018). Each sequence was assigned the best match at a probability better than 90%. Taxonomic information was collected from the Entrez Taxonomy database.

Sequences were clustered with CLANS (Frickey and Lupas 2004) based on the p-value of their BLASTp pairwise comparison, computed using the BLOSUM62 scoring matrix. Clustering of the entire set was preformed until equilibrium at a p-value of 10^−5^ and superclusters identified manually based on the name of the corresponding query sequences and the ECOD domains assigned. To identify subclusters and internal connections, the sequences in the VCBS supercluster, including and excluding the PQQ/RGL11 sequences, were re-clustered at p-values of 10^−18^ (Fig. 1b) and 10^−20^, respectively (Fig. S2a).

In order to evaluate the domain environments of the β-propellers in each subcluster, their parent full-length proteins were collected and binned by size, with a step of 100 residues. A representative for each bin was collected and domains annotated iteratively with HHsearch as above. A maximum of 4 iterations were carried out, where sequence regions not yet mapped to a domain were searched individually. Only the best matches at a probability better than 70% and larger than 40 residues were considered. Signal peptide prediction was carried out with Phobius (Käll et al. 2004).

For HMM comparisons, the sequences composing the clusters and subclusters depicted in figure 1 were used. For each group, the sequences were aligned with MUSCLE (Edgar 2004) and the alignment trimmed with trimAl (Capella-Gutierrez et al. 2009), removing columns where >25% of the positions were a gap (gap score of 0.75) and sequences that only overlapped with less than half of the columns populated by 80% or more of the other sequences. HMM profiles were built with HHmake and aligned with HHalign (Söding 2005), using default parameters without secondary structure scoring. Structural alignments were carried out with TM-align (Zhang and Skolnick 2005).

## Acknowledgements

We thank Felipe Merino for stimulating discussions.

## Funding

This work was supported by ‘Life’-Grant 94810 from the Volkswagenstiftung and by institutional funds of the Max Planck Society.

## Notes

### Competing Interest Statement

The authors have declared no competing interest.

