## Supplementary material for "The VCBS superfamily, a diverse group of β-propellers that includes tachylectin and integrins": S

### SUPPLEMENTARY FIGURES

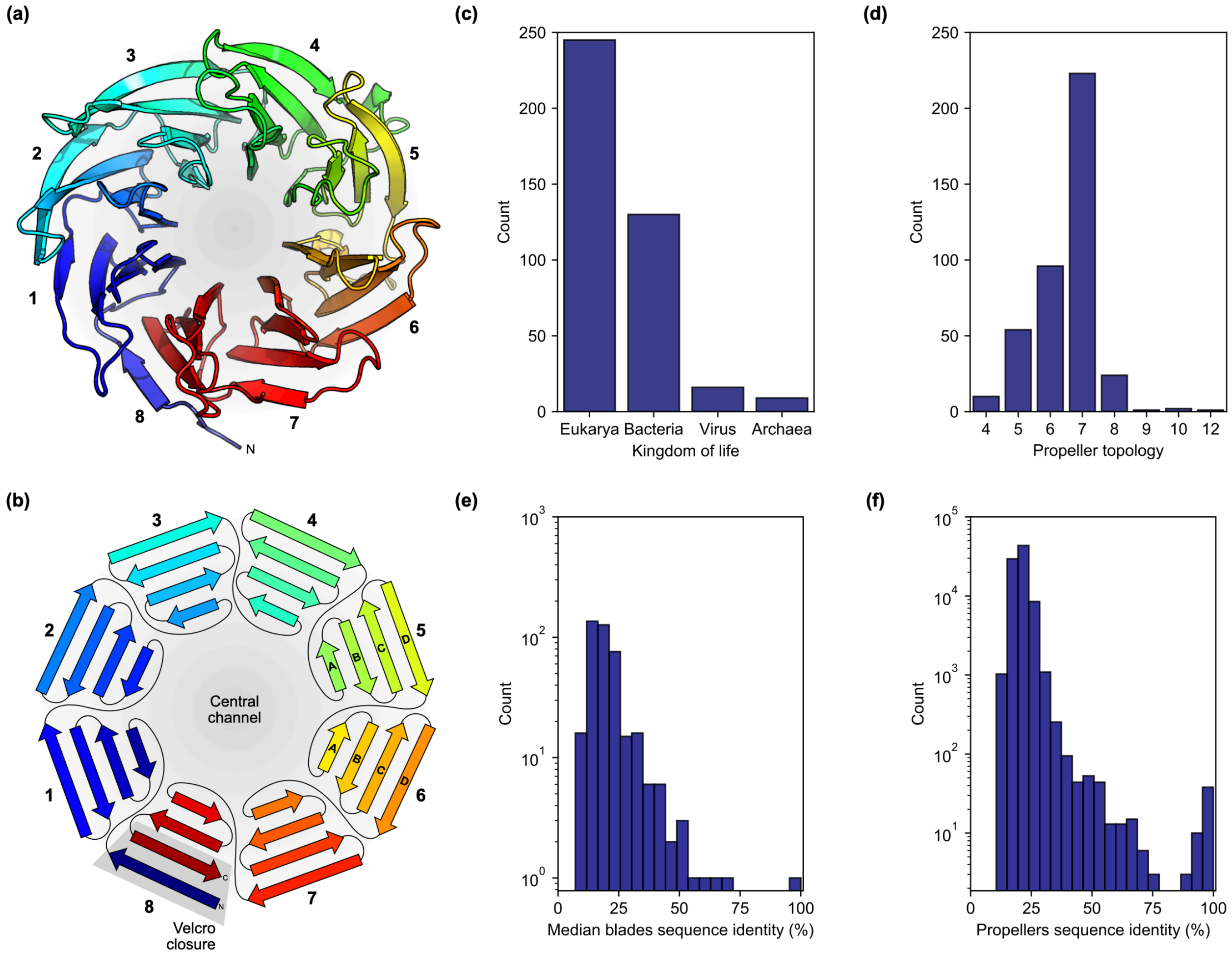


**Figure S1.** General features of the β-propeller fold and its representation in structural databases. (a) 3D structure of a β-propeller, exemplified by the crystallographic model of yeast ribosome assembly protein SQT1 (PDBID: 4ZOV_A), an 8-bladed member from the WD40 supercluster. (b) 2D fold topology of the fold depicted in (a), highlighting the different blades, the A-to-D naming of their constituent β-strands and the characteristic 'velcro-closure'. (c) Taxonomic distribution, (d) number of blades distribution (topology), (e) median pairwise sequence identity between blades within the same β-propeller, and (f) pairwise sequence identity between all β-propeller domains on the Evolutionary Classification of Protein Domains (ECOD) database (Cheng et al. 2014) filtered to a maximum sequence identity of 70% as of January 2020. For computing pairwise sequence identities, sequences were aligned with MUSCLE (Edgar 2004) and only the aligned regions considered.

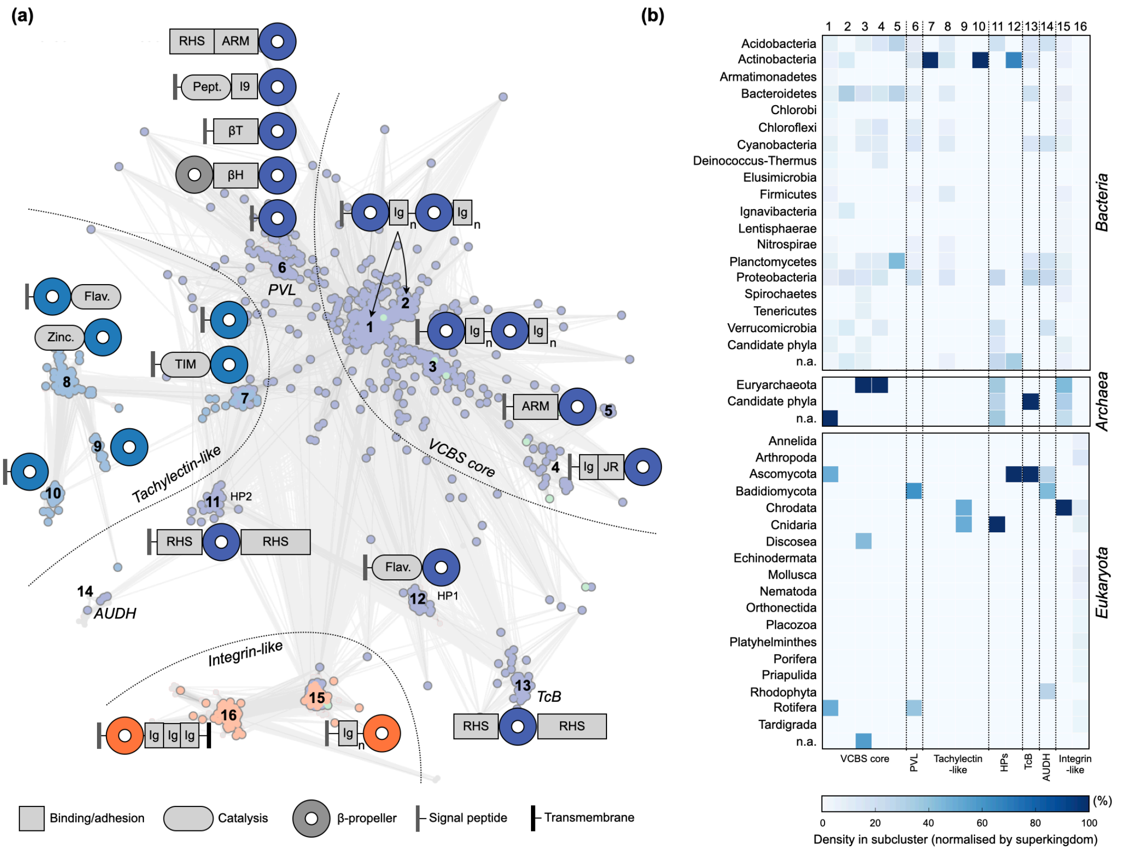


**Figure S2.** Domain environment and taxonomic distribution of β-propellers belonging to the VCBS superfamily. (a) Cluster map of the 1607 β-propeller sequences in the VCBS supercluster in figure 1, excluding those from the PQQ and RGL11 clusters. Clustering was carried out in 2D until equilibrium with CLANS (Frickey and Lupas 2004) at a p-value of $10^{-20}$. Connections are shown at a BLASTp p-value of $10^{-10}$. Example domain compositions of full-length proteins for representatives from different subclusters (identified at a p-value of $10^{-20}$) are shown next to the corresponding region. The architectures shown represent the consensus of the annotations of representatives of a given subcluster; when a consensus was not possible to obtain, examples of the diverse architectures found are shown. Disks denote β-propeller domains, rectangles domains involved in biomolecular interactions and adhesion, and rounded rectangles those involved in catalysis. Ig: immunoglobulin-like; JR: jelly-roll like; ARM: armadillo repeats; RHS: rearrangement hotspot-like repeats; I9: inhibitor I9-like; βT: β-trefoil lectin-like; βH: β-helix like; Pept.: peptidase-like; Flav.: flavodoxin-like; P-loop: P-loop containing ATPase-like; TIM: TIM-barrel like; Zinc.: Zincin-like. A subscript *n* denotes a variable number of units. Signal peptides and transmembrane segments identified are highlighted. HP stands for ‘hypothetical β-propeller’. (b) Taxonomic distribution of the different superclusters, separated by superkingdom. Numbers in the x-axis represent the different subclusters in (a). The darker the cell, the more frequent a given phylum is in a given subcluster. Frequencies are normalised by the total counts of individual superkingdoms in a given subcluster.

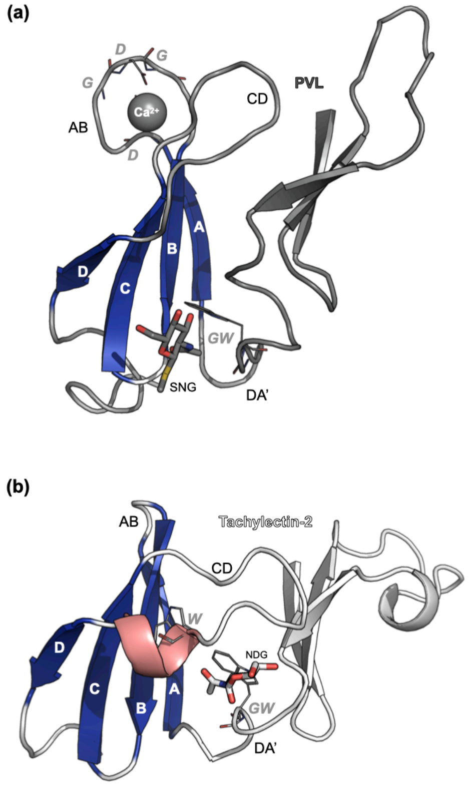


**Figure S3.** Sugar-binding pockets in (a) fungal PVL (pdbID: 2BWM_A) and (b) tachylectin-2 (pdbID: 1TL2_A) β-propellers. The blades in figure 3 and the ones right preceding them are shown, accordingly coloured. The residues in conserved sequence motifs are shown in lines and highlighted, as well as the ligands they interact with. SNG: methyl 2-acetamido-2-deoxy-1-seleno-beta-D-glucopyranoside; NDG: 2-acetamido-2-deoxy-alpha-D-glucopyranose.

### SUPPLEMENTARY TABLES

**Table S1.** Input queries for sequence searches. The ECOD Domain ID is given, as well as the name of the source protein, the domain ECOD topology (t-name) and family (f-name) names, and the supercluster in figure 1 to which they belong to are given.

| Domain ID | Protein name | ECOD t-name | ECOD f-name | Supercluster |
|---|---|---|---|---|
| e5nniA2 | Sortilin | 10-bladed | Sortilin-Vps10 | Asp-Box |
| e2w38A1 | Sialidase | 6-bladed | BNR-2 | Asp-Box |
| e2zuyA1 | YesX | 8-bladed | RGL11-C | VCBS |
| e2bwmA1 | PVL | 7-bladed | VCBS | VCBS |
| e4a7yA1 | AUDH | 7-bladed | VCBS-like | VCBS |
| e4iglA2 | YenB/TcB | Propeller domain in ABC toxin B component | VCBS | VCBS |
| e1tl2A1 | Tachylectin-2 | 5-bladed | Tachylectin | VCBS |
| e1tyeA1 | α-integrin | 7-bladed | FG-GAP-1st | VCBS |
| e1rwiA1 | PknD | 6-bladed | SGL-4 | WD40 |
| e1ijqA1 | LDL receptor | 6-bladed | SGL-5 | WD40 |
| e1tbgA1 | Transducin/G-protein | 7-bladed | ANAPC4-WD40-3 | WD40 |
| e1l0qA2 | Surface layer protein | 7-bladed | Lactonase-1 | WD40 |
| e3p1lA1 | BamB | 7-bladed | PQQ-2 | VCBS/PQQ |
